# Single-molecule mass measurements uncover shifting RNA interactions during condensate phase transitions

**DOI:** 10.64898/2026.07.27.740926

**Authors:** Axel Leppert, Jesper Shiapan, Irena Papageorgiou, Qi Ying Neo, Cecilia Mörman, Hannah Osterholz, Petra Mészáros, Max F. Hantke, Dilraj Lama, Ali Miserez, Axel Abelein, Michael Landreh

## Abstract

RNA interactions are a key contributor to the formation and disassembly of intracellular protein condensates. Although some proteins utilize specific RNA-binding domains, these processes can also be mediated by charge interactions with intrinsically disordered regions. Due to the dynamic nature of these systems, investigating the underlying specificity and stoichiometry remains challenging. Here, we demonstrate that single-molecule mass measurements with mass photometry can capture RNA-protein interactions in phase-separated protein systems. Using the approach to investigate RNA-mediated phase shifts of tau condensates, we find that increasing the RNA concentration, which promotes phase re-entry, results in RNA-mediated tau multimerization, where each tau monomer binds a linear RNA sequence of approximately 30 nucleotides. Solution NMR and native mass spectrometry confirm the formation of stable complexes between RNA and the basic proline-rich and repeat domains of tau, which have a net charge of -29. Our findings demonstrate that mass photometry can distinguish between charge neutralization, which drives coacervation, and complex formation, which mediates phase re-entry, making it a highly complementary tool for the study of RNA-mediated phase separation.

## Introduction

Protein–RNA interactions are a key component of cellular organization and regulation.^1^ Many RNA-related processes, such as transcription, translation, and RNA processing, are compartmentalized into biomolecular condensates, membrane-less assemblies that can be dynamically assembled and disassembled in the cell.^2^ Often, RNA is not only a biochemical substrate but actively regulates the condensate formation.^1,2^ By engaging in multivalent interactions with proteins, RNAs can promote phase separation and stabilize condensates, while changes in RNA concentration or sequence can also trigger condensate dissolution or remodeling.^3,4^

RNA is recruited into condensates by RNA-binding proteins that undergo liquid-liquid phase separation (LLPS) through multivalent interactions between disordered domains. Different types of RNA interactions are illustrated by TDP-43, Fused in Sarcoma (FUS) and microtubule-associated tau protein, all of which form condensates *in vitro* alone or in the presence of RNA.^5–7^ TDP-43 contains tandem folded RNA recognition motifs (RRM) that mediate preferential binding of GU- and UG-rich RNAs.^7,8^ FUS interacts with RNA through both a canonical RNA-binding domain and its arginine-rich N-terminal region, enabling both specific GGU motif recognition and non-specific RNA binding.^9^ Tau binds RNA solely through its disordered microtubule binding domain, which is rich in basic residues, with no specificity except a preference for single strands.^10–12^ At a 1:1 charge balance, where there is a positively charged sidechain (R, K, H) for each negatively charged backbone phosphate, the negatively charged RNA can neutralize the positive net charge of tau, leading to complex coacervation.^11,13^ Increasing the RNA concentration beyond charge balance inhibits condensate formation.^11^ The formation of overcharged complexes has been proposed as the basis for disassembly, but their specific molecular interactions remain unclear.^3,4^ Understanding the underlying mechanisms of RNA-driven phase separation and condensate dissolution is of particular interest for the development of drug delivery systems. Phase-separating peptides rich in aromatic and/or basic residues can encapsulate mRNAs into coacervates and release them following external cues.^14–16^ Therefore, monitoring interactions between cargo and vehicle under assembly and disassembly conditions at the molecular level can offer crucial insights for the design of targeted delivery systems.^17,18^

Together, these examples demonstrate the range of protein architectures that mediate RNA interactions in condensates. Specific RRM-RNA complexes, including FUS and TDP-43, have been resolved with X-ray crystallography.^9,19^ Furthermore, cryo-EM structures of fibrils formed by the basic domains of tau and FUS indicate a direct interaction with RNA.^20,21^ However, despite their importance for the regulation of condensate formation, the structural features that govern interactions with basic, disordered regions remain poorly understood. This knowledge gap is largely due to the technical challenge to determine the underlying binding stoichiometries and interaction preferences in such dynamic systems.

Here, we demonstrate that mass photometry (MP), which enables single-molecule mass measurements of proteins, DNA, and, most recently, RNA,^22–24^ reliably captures RNA-mediated oligomerization of disordered protein domains. With MP, we can follow mRNA cargo loading into engineered peptide coacervates developed for intracellular drug delivery. Using a newly developed calibration strategy, we monitor the formation of protein-RNA complexes during RNA-dependent tau phase transitions. We find that phase re-entry of tau condensates is driven by the formation of overcharged, stable complexes in which each tau molecule occupies an approximately 30-nucleotide segment of RNA through interactions with its basic region. These complexes are not observed under LLPS conditions. Our results establish mass photometry as a viable approach to unravel the heterogeneity of non-specific protein-RNA interactions and their role in condensate formation and dissolution.

## Results

### MP captures interactions between RNA and disordered proteins

MP is routinely applied to detect homotypic interactions of proteins or nucleic acids, and, most recently, interactions between RNA and antibodies.^23–26^ We thus reasoned whether it could detect heterotypic interactions between the negatively charged phosphates in the RNA backbone and basic sidechains in disordered protein domains that drive coacervate formation. To test this possibility, we turned to designed disordered proteins that contain a spider silk-derived folded N-terminal domain (NT*) with a C-terminal tail composed of 14 repeats of GSGAP, GSGAE, or GSGAK, recapitulating features of native low-complexity domains (***Table S1, Figure 1a***).^27^ Proteins were incubated with or without a 50-nucleotide Poly-A RNA and then measured by MP. Importantly, the proteins and RNA alone are below the *ca*. 30 kDa-detection limit of the mass photometer^28^ (22 kDa and 16 kDa, respectively), whereas complexes between protein and RNA are above the limit (37 and 59 kDa, respectively) (***Figure 1b***). Measurements were conducted using lysine-coated slides to facilitate detection of negatively charged RNA.^23^ As expected, no signal was detected when proteins and RNA were measured separately, as indicated by the comparable intensities of the positive and negative mass binding events in the histograms,^28^ which match those seen in buffer-only histograms (***Figure S1***). After incubation of proteins and RNA together at a 1:1 ratio, we observed a strong signal for the basic protein variant (GSGAK), which is significantly larger than the anti-binding peak. The broad mass of 45 ± 11 kDa is compatible with a complex between one RNA and one or possibly two GSGAK proteins. Addition of RNA to the neutral GSGAP and the acidic GSGAE variants, on the other hand, caused no change in intensity of the binding and anti-binding peaks (***Figure 1c***). The presence of high sodium chloride concentrations reduced the MP signal for the RNA-protein complex, as expected for a charge-based interaction (***Figure S1***). Since exact mass determination at the detection limit of MP is challenging, we turned to native mass spectrometry (nMS) to confirm the interaction between the GSGAK protein and the RNA. At the same buffer conditions as for the MP measurements, the resulting spectra show a pronounced 1:1 interaction between protein and RNA with low charge states expected for protein-nucleic acid complexes (***Figure S1***).^29^

**Figure 1.**
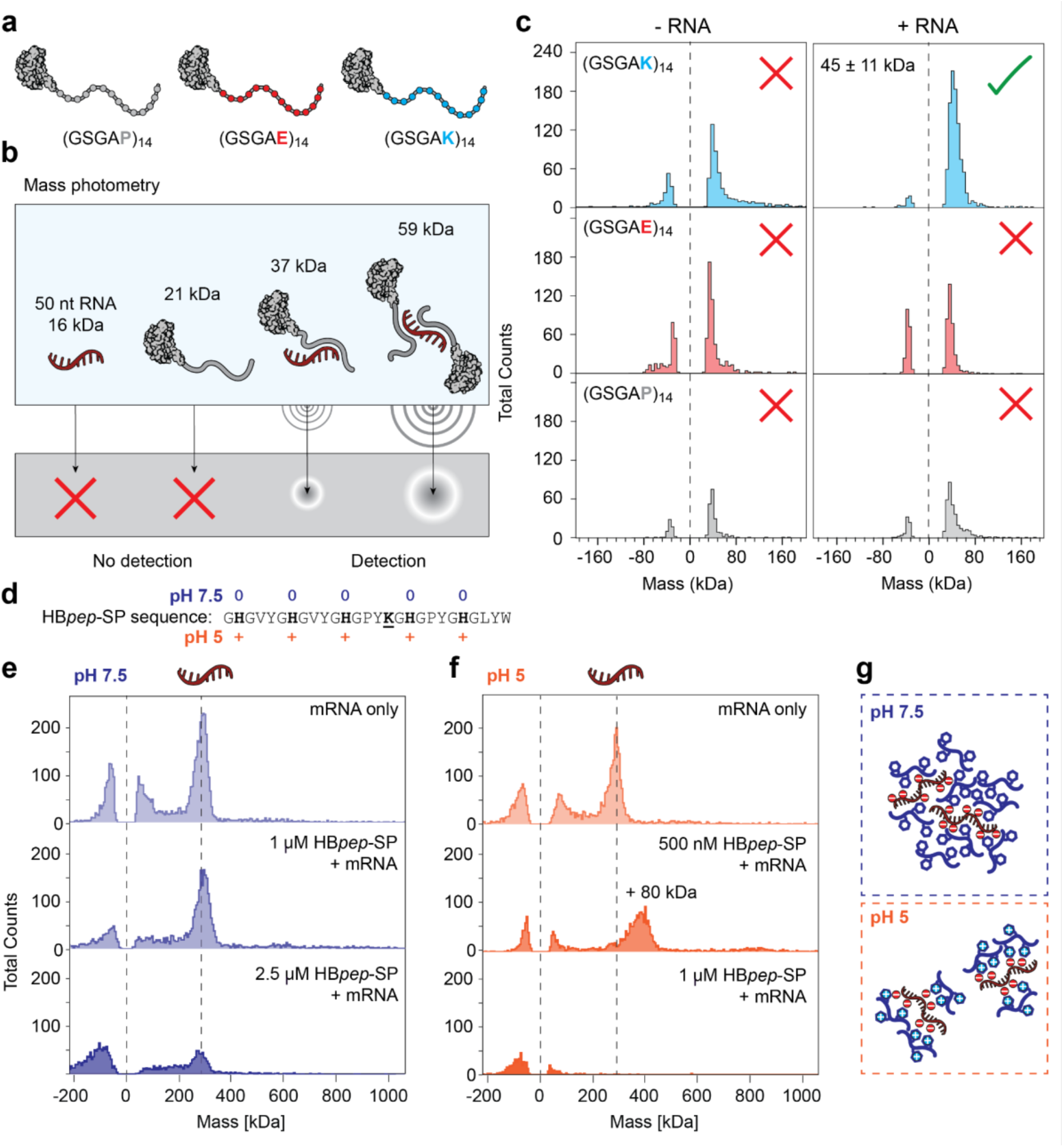
A mass photometry assay for protein-RNA interactions. (a) Proteins with model low-complexity domains are composed of a spider silk-derived NT* domain (grey) and 14 repeats of GSGAP, GSGAE, or GSGAK, representing neutral, acidic, and basic disordered sequences. (b) Principle of the RNA binding assay. Individually, proteins and RNA masses are below the detection limit, whereas each possible complex has a mass above the limit, facilitating detection. (c) MP histograms of each model protein incubated with a 3:1 charge ratio of 50-nucleotide poly-A RNA show a detectable complex with a broad mass distribution between the basic protein variant (GSGAK)_14_ and the RNA. (d) Sequence of HBpep-SP with charges at pH 7.5 and 5 indicated. The single lysine residue (underlined) is modified with a redox-cleavable moiety. (e) MP histograms of EGFP mRNA in the presence of 0 - 2.5 μM HBpep-SP recorded at pH 7.5 show no change in mRNA mass and reduced signal intensity at the highest peptide concentration. (f) At pH 5, the presence of HBpep-SP at a concentration of 500 nM leads to a mass shift of approximately 80 kDa for the mRNA. Higher peptide concentrations cause loss of signal. (g) pH-dependent change in interactions between HBpep-SP and RNA. At pH 7.5, the neutral peptide is able to undergo self-assembly and recruit RNA. At pH 5, the positive charge of the peptide enables strong binding to negatively charged RNA but inhibits self-assembly.

These findings raise the possibility to probe assembly mechanisms of peptide-RNA coacervates. As a test case, we selected the squid beak-derived HB*pep*-SP system capable of encapsulation and intracellular delivery of mRNAs.^15^ The histidine-rich HB*pep*-SP spontaneously forms coacervates at neutral pH, whereas acidic pH inhibits LLPS (***Figure 1d***).^30^ Cargo molecules that are present during peptide assembly are incorporated into the coacervates.^15^ We reasoned that the ability of MP to detect protein-RNA complexes could be used to probe the interactions between HB*pep*-SP and mRNA under LLPS and non-LLPS conditions. For this purpose, we measured the mass of an mRNA encoding EGFP together with increasing amounts of HB*pep*-SP at pH 7.5 (***Figure 1e***). At pH 7.5, when the peptide is neutral, the mass of the mRNA remains unchanged in the presence of 10 nM - 1 μM peptide. At 2.5 μM peptide, the RNA signal intensity is reduced, and above 5 μM peptide, the RNA could no longer be detected (***Figure 1e, Figure S1***). Next, we repeated the experiment at pH 5, where the histidines carry a positive charge. Here, the presence of 500 nM HB*pep*-SP shifts the mass of the mRNA by approximately 80 kDa, corresponding to binding of *ca*. 20-25 peptide monomers. Increasing the peptide concentration to 1 μM leads to loss of signal, which could be due to the formation of larger assemblies (***Figure 1f, Figure S1***). These data are consistent with a shift in RNA interactions: at pH 7.5, neutral HB*pep*-SP can form self-assemblies and recruit RNA, likely through π-π stacking between the bases and histidine and tyrosine residues.^31,32^ At pH 5, protonated HB*pep*-SP does not self-assemble, but is able to bind with high affinity to the negatively charged RNA, forming small complexes that can be detected by MP (***Figure 1g***). In summary, MP is suitable for the detection of different interaction modes between positively charged disordered domains and negatively charged RNA.

### Phase re-entry is mediated by binding of 30 nt RNA to each tau monomer

Since MP can detect protein-RNA complexes, we sought to apply this approach to follow RNA-mediated phase transitions of a condensate-forming protein. The disordered tau protein associates with RNA in cells and undergoes phase separation and phase re-entry in an RNA concentration-dependent manner.^11^ Tau exhibits a complex phase behaviour with competing crowding- and charge-driven processes involving the negatively charged projection- and C-terminal domains, as well as the positively charged proline-rich and microtubule binding domains.^5,12^ We therefore first sought to establish experimental conditions that allow us to observe RNA-mediated coacervation of full-length tau Reducing the concentration of ammonium acetate (AmAc), a standard buffer for mass measurement with mass spectrometry (MS), from 100 to 2.5 mM leads to homotypic LLPS (***Figure S2***).^33^ At 25 mM, we observe few individual condensates, suggesting that the protein is at the onset of droplet formation. We then added polydisperse poly-A RNA with an average length of 300 nucleotides (Figure S2). Since the net charge of tau at pH 8.0 is 2+, addition of one 300 nt RNA per 150 tau monomers corresponds to a 1:1 charge ratio. At 25 mM AmAc, 1:1 charge matching between tau and RNA resulted in pronounced phase separation. Fluorescence microscopy confirmed the presence of tau and RNA in the droplets in all cases (***Figure 2a***). Increasing the RNA concentration to a 3-times molar excess over tau (300 nt RNA per tau monomer) caused complete dissolution of the droplets under all buffer conditions (***Figure 2a, Figure S2***). We therefore chose 25 mM AmAc as the basis for monitoring RNA-mediated transitions between one-phase, two-phase, and re-entrant one-phase systems.

**Figure 2.**
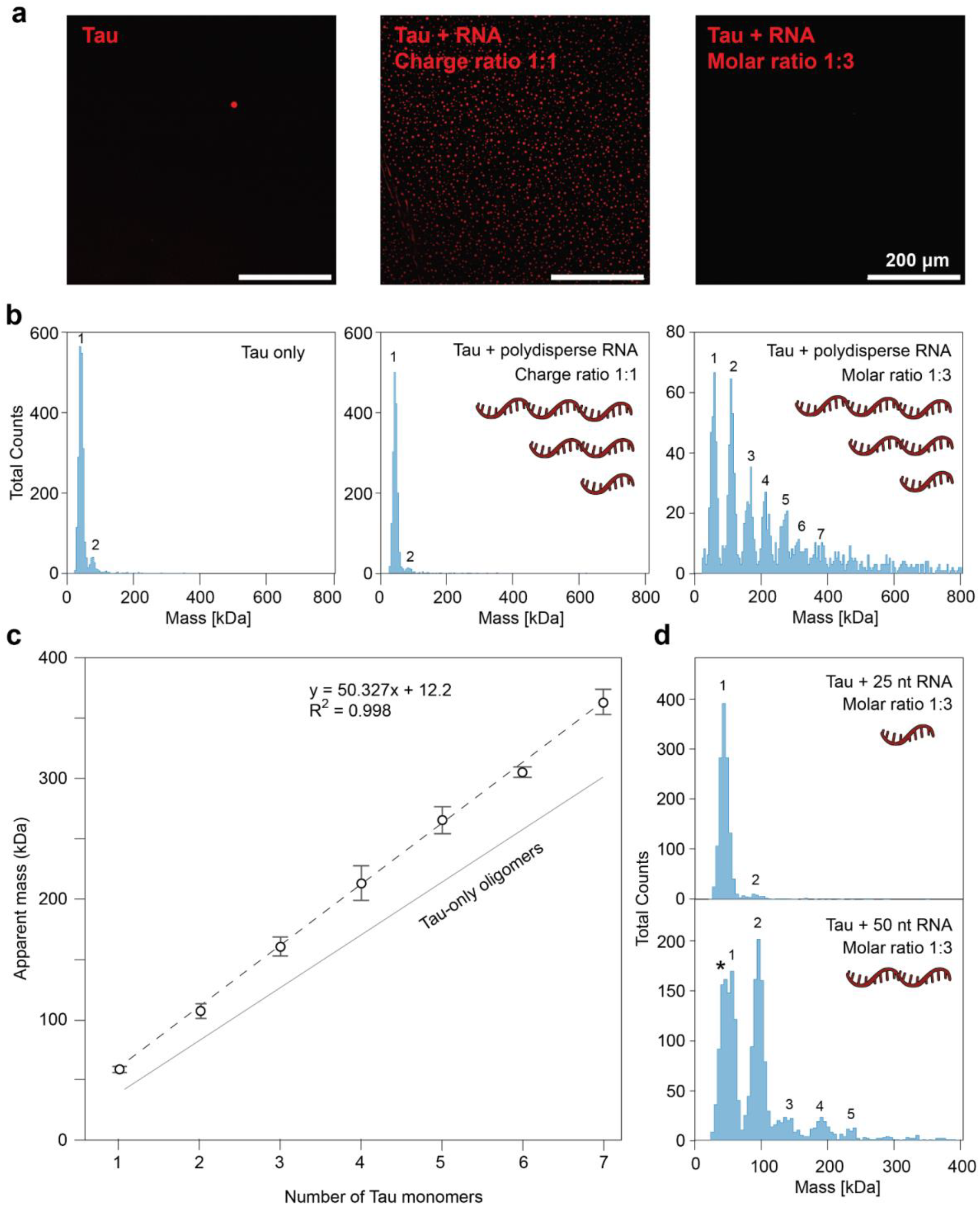
Phase re-entry is accompanied by binding of 30 nt RNA per tau monomer. (a) Fluorescence microscopy of 15 μM tau (+1% Atto655 labeled tau, red) in 25 mM AmAc without RNA (left), with polydisperse poly-A RNA (shows as three RNA molecules) at a charge ratio of 1:1 (100 nM RNA with an average length of 300 nucleotides, middle), and at molar ratio of 1:3 (5 μM RNA, right). Coacervates form at a 1:1 charge ratio. Scale bars are 200 μm. (b) MP histograms of 9 nM tau in 25 mM AmAc alone (left), with 60 pM polydisperse poly-A RNA (charge ratio 1:1, middle) and with 3 nM RNA (molar ratio 1:3, right). The number of tau monomers is indicated above each peak. (c) Plot of the observed tau-RNA complex masses as a function of oligomeric state. Linear fit shows an average mass of 50.3 kDa per tau monomer. Data are shown as average and standard deviation (n = 3). The theoretical masses for tau-only oligomers are indicated by a solid line. (d) MP histograms of tau with 25- and 50-nucleotide poly-A RNA at a 1:3 molar ratio. Only the 50-nucleotide RNA induces tau dimerization. * indicates a shoulder peak corresponding in mass to tau with no RNA.

Having established conditions for RNA-mediated phase transitions of tau, we investigated their interactions using MP. Importantly, MP requires low nanomolar analyte concentrations. Though being below the threshold for macroscopic droplet formation, these conditions allow the observation of nanoscale clusters that represent early steps in phase separation.^34^ MP histograms of tau in the presence of 25 mM AmAc show a strong monomer signal with the expected mass of 42 kDa, as well as a minor dimer population, in line with previous studies **(Figure 2b)**.^33^ Addition of polydisperse poly-A RNA at a charge ratio of 1:1 does not cause notable changes in the histograms. The data suggest that RNA does not have a measurable effect on tau oligomerization at the sub-stoichiometric RNA concentrations required to form droplets **(Figure 2b, c)**. Next, we analyzed tau and RNA at a molar ratio of 1:3, where no coacervates are present. Strikingly, RNA caused the appearance of well-defined oligomers ranging from 50 to > 500 kDa. The oligomer signals are evenly spaced, which indicates multimers containing between one and seven tau molecules. Plotting the masses of the oligomers as a function of the oligomeric state reveals that each subunit has an apparent molecular weight of 50.3 kDa, 7 kDa above the mass of tau (***Figure 2c***). The same effect was observed for 2.5 and 25 mM AmAc, but not for 100 mM AmAc, which appeared to reduce tau-RNA interactions (***Figure S3***). We speculated that the mass increase is due to multivalent interactions with RNA strands that connect the tau monomers. However, since single-strand RNA has a different refractive index than proteins, the mass of the RNA component could not be determined together with the tau oligomers. To correct for the difference, we measured a series of RNA molecules with known masses using the standard protein calibration (***Figure S3***). We found that the protein calibration under-estimates the mass of the RNA by 25%. From this trend, we can derive a correction factor of 1.34. By multiplying this factor with the RNA mass measured with a protein calibration, we can accurately determine the mass of the RNA component in the oligomers to be 10.1 kDa per tau monomer. Assuming an average weight of 340 Da per nucleotide, this value corresponds to approximately 30 bases. In addition, we detect an additional mass of 16 kDa that is constant regardless of oligomer size, reflecting potentially a minimum length requirement for the first bound RNA.

The data suggest that at a molar ratio of 1:3, a single RNA can bind multiple tau molecules, each occupying an average of 30 bases. To test this hypothesis, we recorded MP histograms of tau with synthetic poly-A RNAs containing either 25 or 50 bases. With the 25-base RNA, no tau oligomers were observed, whereas the 50-base RNA led to the formation of tau monomers and dimers, as well as a small amount of trimers and tetramers (***Figure 2d***). For these oligomers, we determined an additional mass of *ca*. 13 kDa, which did not increase with oligomer size (***Figure S3***). From these data we conclude that 50 nucleotides, but not 25, bind tau with enough overhang to allow binding of a second tau molecule. These data are in good agreement with the observed occupation of 30 nucleotides per tau monomer.

### RNA binds in extended conformation to the microtubule-binding region of tau

The surprisingly well-defined requirement of an average of 30 nucleotides per tau molecule raises the question of how such a defined interaction length is encoded in the intrinsically disordered tau sequence. As the first question, we asked whether the requirement for 30 nucleotides represents the spacing between two tau monomers bound to the same RNA, or forms a single interaction interface for one tau monomer. We therefore measured binding between poly-A RNA and tau at a molar ratio of 1:3 using nMS. Collisional activation was required to obtain well-resolved mass spectra, suggesting cluster formation from which stable protein-RNA complexes can be released by fragmentation (***Figure S4***). The resulting spectra show that tau monomers preferentially bind up to two 25-nucleotide RNAs but only a single 50-nucleotide RNA (***Figure 3a***). The data suggest that one tau can accommodate more than 25 bases, whereas 50 bases completely saturate binding, in good agreement with the finding from MP that each tau occupies 30-nucleotide stretches on longer RNAs. Interestingly, we do not observe a 2:1 complex between tau and the 50-nucleotide RNA seen in MP (***Figure 2d***). A possible reason could be the collapse of the nucleic acid in the gas phase, which may reduce the available interaction surface.^29^ This difference highlights the ability of complementary mass measurement approaches to capture different interaction modes.

Since the nMS data indicates that the length requirement stems from lateral binding of the RNA along the protein, we sought to identify the tau region that mediates the interaction. Previous NMR studies have identified pronounced interactions between folded tRNAs and the basic regions of tau, which were predominantly mediated by arginines in the proline-rich domain.^35^ Expanding on these findings, we recorded 2D NMR ^1^H-^15^ N-HSQC spectra to determine which residues exhibit modulation of signal intensity and chemical shift changes (CSC) upon titration with polydisperse PolyA RNA (***Figure 3b,c, Figure S5***). From 0.2 - 0.7 μM Poly-A RNA additions onto 128 μM ^15^ N-labeled tau, which correspond to concentrations below RNA-mediated tau phase separation, only minor changes in the spectra were observed (***Figure S5***). The largest perturbations were found for positively charged residues. Upon increasing Poly-A concentrations, approaching RNA-mediated phase separation at approximately 1.9 μM, larger intensity loss and CSC were detected. The most pronounced changes with 40-90% loss of signal intensity and significant CSC were observed for residues 225-260 and 290-350, corresponding to the P2/R1 and R2/R3 tau regions in the basic domain (***Figure 3b, Figure S5***). Upon additional PolyA RNA titrations steps, previous perturbations were enhanced and at the final concentration of 42.7 μM Poly-A RNA, corresponding to a molar ratio of three tau to one RNA, the P1 region was also affected. Interestingly, the proline-rich and repeat domains are predicted to exhibit a net charge of 29+, which is in good agreement with binding of approximately 30 negatively charged nucleotides (***Figure 3c***).

**Figure 3.**
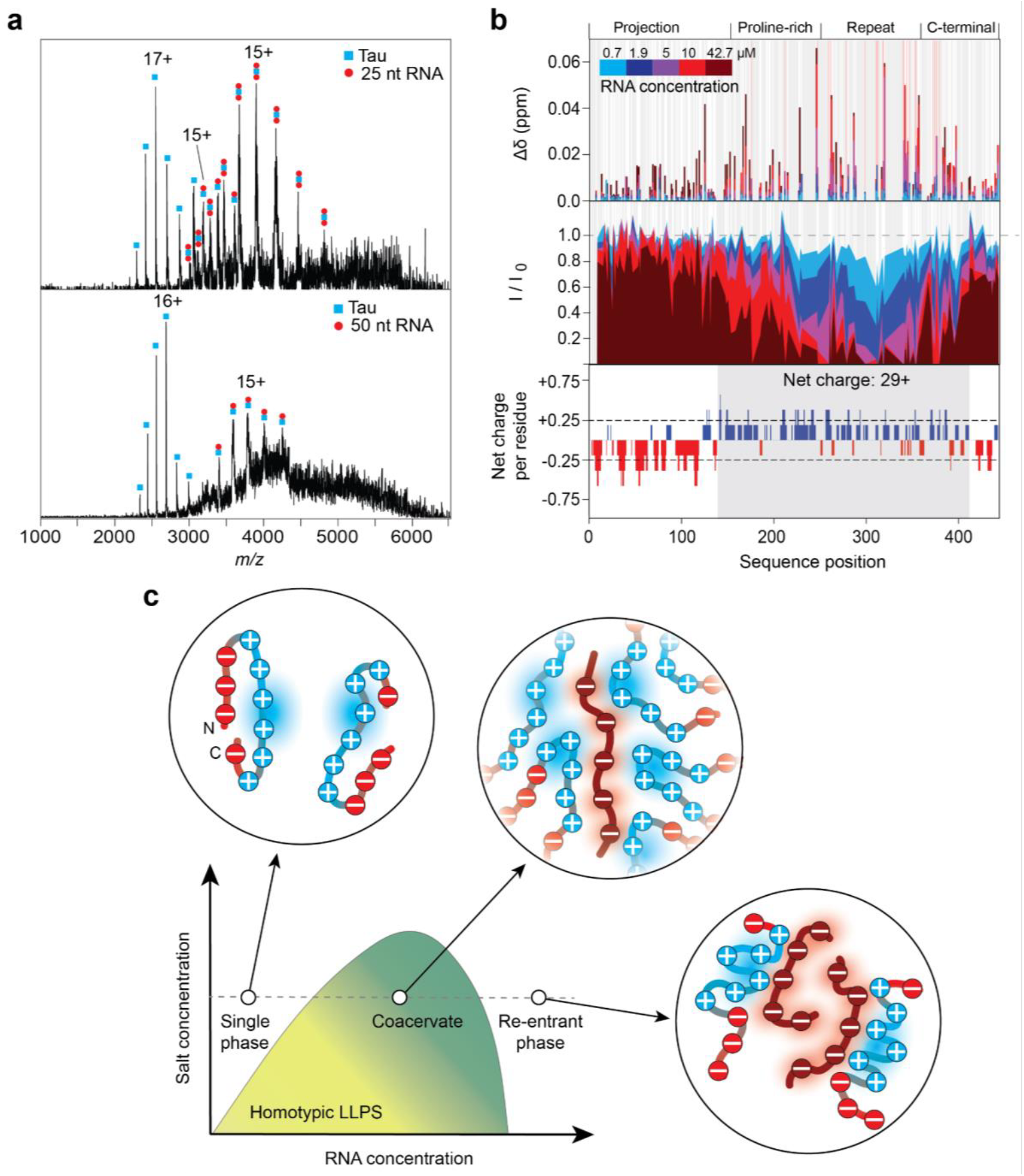
Different-length RNAs bind to the tau repeat domain. Figure 3. RNA binds in an extended state to the tau repeat domain. (a) Native mass spectra of tau in the presence of poly-A RNA show non-covalent complexes. One tau molecule can form a stable complex with two 25-(top) or one 50-nucleotide poly-A RNA (bottom). (b) Top panel: 2D NMR ^1^H-^15^N HSQC spectroscopy of 128 μM ^15^ N-labeled tau and titration with polydisperse poly-A RNA. Top: Bar plot of quantified CSCs at each titration step for each assigned residue, with grey background bars representing residues unable to be assigned, and red background bars representing residues which had signal losses enough for the cross-peak to disappear from the spectra. Middle panel: relative intensity, with grey background bars representing unassigned residues. Bottom panel: Plotting the net charge per residue using CIDER^36^ shows a correlation between basic domains and RNA-induced signal attenuation and CSCs. (c) A shift from indirect to direct RNA interactions mediates phase transitions of tau. The relative positions of the three conditions evaluated here (no RNA, charge ratio 1:1, and molar ratio 1:3) are indicated in the schematic tau-RNA phase diagram. Without RNA, and at moderate salt conditions, tau is partially compact with a positive net charge that prevents coacervation. At 1:1 charge ratio, transient contacts with RNA neutralize the positive charge, and tau phase-separates into coacervates with RNA without forming stable complexes. At 1:3 molar ratio, the repeat regions of multiple tau molecules are stably bound by RNA, which sequesters sufficient amounts of tau into highly negatively charged complexes to drive re-entry of the system into a single phase.

To confirm this hypothesis, we performed all-atom MD simulations of tau residues 280-360 with 30 nucleotide poly-A RNA and quantified the per-residue contacts. As expected, we find strong interactions with the basic lysine residues in the sequence, with particularly K321 and K331 showing near-constant interactions with the RNA backbone (***Figure S4***). In summary, we observe mild CSPs near the center of the repeat domain at low RNA concentrations, which intensify at high concentrations and cover the entire repeat and proline-rich domains. Considering the stoichiometries observed by MP and nMS, we conclude that low amounts of RNA engage in transient or weak contacts with the center of the repeat region. As the amount of RNA increases, these contacts are replaced by the formation of stable complexes that cover the length of the basic domains.

## Discussion

MP is widely used to determine the masses of proteins, DNA, and RNA. However, protein-nucleic acid interactions have not previously been studied due to different refractive indices of their components, which results in inaccurate mass measurements. We demonstrate that this issue can be overcome for complexes where the relative nucleic acid content can be estimated. We determine a scaling factor of 1.34 which can be applied to determine the molecular weight of the bound nucleic acid.

As a direct application of the approach, we monitored interactions between tau and RNA during coacervate formation and phase re-entry. The ability of RNA to modulate tau phase transitions in a charge ratio-dependent manner is well-established,^11,12,12,13^ but the molecular interactions that drive these processes have been challenging to study. MP reveals that at a 1:1 charge ratio, no direct interactions between the components can be detected. Increasing the RNA concentration 3-fold leads to the formation of “beads on a string”-type complexes in which RNAs of varying lengths add tau molecule for every 30 nucleotides. We speculate that the use of lysine-coated MP slides may bias the measurements towards negatively charged species, yet no complexes are detected at a 1:1 charge ratio, which implies that they only form once the RNA concentration is increased (***Figure 3c***).

MP thus confirms that tau-RNA coacervation is driven by charge matching, an indirect interaction where each RNA neutralizes a large number of tau molecules. Increasing the RNA concentration induces a sudden shift from indirect to direct interactions. Binding of multiple tau molecules to long RNAs thus sequesters the protein into negatively charged complexes, which lowers tau solution concentration and increases Coulombic repulsion. Both factors likely drive coacervate disassembly and phase re-entry (***Figure 3c***).

Lastly, we considered how such a shift in interactions could be mediated. The NMR data indicate that contacts at low RNA concentrations involve predominantly the tau repeat region, which harbours 18 lysines but contains only a single arginine. Interactions at high RNA concentrations cover the proline-rich region, which contains 8 lysines and 8 arginines. Lysine interacts weakly with the phosphate backbone, whereas arginine preferentially forms more stable contacts.^35,37^ Lysine interactions contribute to tau-RNA coacervate formation, which is inhibited by lysine acetylation.^35^ We speculate that our use of polydisperse poly-A RNAs favors more heterogeneous contacts, which nevertheless supports the critical role of lysines in mediating coacervation with RNA. In solution, tau can adopt a “paperclip” conformation in which the basic domains interact with the negatively charged N- and C-termini.^38,39^ We speculate that different affinities of the repeat domain and proline-rich domains for RNA, as well as competition between terminal domains and RNA for binding to the basic domains of the protein, could account for RNAS-induced phase transitions.

In conclusion, we establish that MP informs about interactions of disordered protein domains with RNA, and use the approach to uncover how a shift from indirect to direct RNA interactions mediates phase transitions of the human tau protein. Beyond providing insights into structure and dynamics of tau-RNA coacervates, we anticipate that this work can serve as a blueprint for detailed investigations into a wide range of heteromolecular condensates.

## Materials and Methods

### Peptide and mRNA synthesis

HB*pep*-SP variant HP was synthesized and purified as described.^30^ The DNA template for EGFP mRNA synthesis was generated by polymerase chain reaction amplification of a custom-designed pcDNA-EGFP plasmid using Takara SeqAmp™ DNA polymerase and gene-specific primers. In vitro transcription (IVT) was performed using the NEB HiScribe™ T7 High Yield RNA Synthesis Kit according to the manufacturer’s protocol. N1-methyl-pseudouridine-5’-triphosphate, CleanCap AG and RNase inhibitor were incorporated in the reaction to improve mRNA stability, capping efficiency and overall transcript yield. The resulting EGFP mRNA was characterized using an RNA TapeStation assay to confirm transcript size and integrity. Poly-A25, Poly-A50 were purchased from GenScript and poly-A polydisperse RNA from Sigma-Aldrich (P9403).

### Production of (GSGAX)_14_ protein variants

All chemicals were purchased from Sigma unless noted otherwise. Plasmids were ordered from GeneScript. NT*-(GSGAE)_14_, and NT*-(GSGAP)_14_, plasmids were transformed into competent *E. coli* BL21 (DE3) cells. Single colonies were inoculated into 20 mL LB medium with ampicillin (100 µg/mL) and grown overnight (37 ºC, 200 rpm). 5 ml start culture were added into 500 mL LB medium containing ampicillin (100 µg/mL), cells were grown until an OD600 of 0.7-0.9 protein and expression was induced by adding 0.5 mM IPTG (cells grown for 4h at 30 ºC, and 200 rpm). The same procedure was used for NT*-(GSGAK)_14_ with the difference that carbenicillin (100 µg/mL) was used instead of ampicillin, expression was induced with 1 mM IPTG, and the cells were grown overnight at 30 °C. Cells were harvested by centrifugation (20 min, 5000 g, 4 ºC) and resuspended in 20 mM Tris, pH 8, with protease inhibitor. After sonication (6 min, 30% amplitude, 2 s on 10 s off) the lysate was centrifuged (20 min, 20 000 g, 4 ºC) and the protein was purified from the filtered (0.22 µm) supernatant by Ni affinity chromatography eluting bound protein with a gradient from 20 mM Tris pH 8 buffer to 20 mM Tris, 500 mM imidazole pH 8 buffer. Collected fractions were dialyzed in 20 mM Tris pH 8 overnight to remove imidazole, and the His-tag was cleaved by adding TEV protease (1:50 w/w) overnight. In a final reverse IMAC, TEV and uncleaved protein were removed and the cleaved protein was collected and concentrated. The same procedure was used for NT*-(GSGAK)_14_ with the difference that from sonication onward, 500 mM NaCl were used in all buffers.

### Production of tau and ^15^ N-labeled tau

Human tau 1-441 (2N4R) (thereafter referred to as tau) was expressed and purified as previously described.^40^ Briefly, the pET15b-Tau recombinant T7 expression plasmid was transformed into *Escherichia coli* BL21(DE3) cells. Expression was induced by 0.4 mM IPTG for 2 h at 37 °C. The cells were harvested followed by sonication of the resuspended pellet in 50 mM sodium phosphate buffer pH 6.5 supplemented with 1 mM EDTA and protease inhibitor cocktail tablet (cOmplete, EDTA-free, Roche). After centrifugation at 4 °C to remove large cell debris, the supernatant was collected and heat-treated followed by centrifugation. To further purify the tau protein, cation-exchange chromatography (HiTrap SP HP, Cytiva) was applied with elution using an increasing NaCl gradient. Fractions were collected and analysed by SDS-PAGE, and pure tau protein fractions were pooled together. NaCl was removed by a desalting column into the buffer of choice, and stored at -20 °C until used. The tau concentrations were determined using absorbance at 280 nm and an extinction coefficient of ε = 7550 M^-1^cm^-1^. For production of ^15^ N-labeled human tau, the same protocol was used, but minimal M9 media and addition of ^15^ NH4Cl were used instead of LB media.

### Tau labeling and sample preparations

Tau was fluorescently labeled after exchanging the protein into a 10 mM sodium phosphate buffer, pH 8.3, using Micro Bio-Spin 6 columns (Bio-Rad). Atto655 NHS ester (Sigma-Aldrich) was freshly dissolved in acetonitrile and added to the protein at a twofold molar excess. The reaction was allowed to proceed for 30 min at room temperature. Excess unreacted dye was subsequently removed by a second round of buffer exchange into a 20 mM sodium phosphate buffer, pH 8 using micro Bio-Spin 6 columns. The degree of labeling was calculated from absorbance measurements at 280 nm and the Atto655 absorption maximum and was approximately 80%. Prior to phase separation experiments, stock solutions of tau and Atto655-labelled tau were buffer exchanged into 25 mM ammonium acetate pH 8 using 7K MWCO Zeba™ Spin Desalting Columns (ThermoFisher Scientific 89877/89878). Lyophilized aliquots of Poly-A25, Poly-A50 (GenScript), or poly-A polydisperse RNA (Sigma-Aldrich; P9403) were reconstituted with RNase free Ultrapure water (ThermoFisher Scientific, 10977015) to concentrations of 100 μM or 200 mg/mL for poly-A polydisperse RNA. Aliquots and samples were prepared using RNase free, low binding tubes (Eppendorf, EP0030108051). Additionally each RNA type was labelled with Cy3 using Mirus bio Label IT® Nucleic Labeling Kit (MIR 3625) following the manufacturers instructions.

### Microscopy

Samples were prepared in non-binding, black, half-area 96-well polystyrene microplates with a transparent bottom (Corning, CLS3881) and imaged at room temperature. Imaging was performed using a Leica STELLARIS 5 confocal microscope operated with LAS X software version 4.6.1.27508. Excitation was provided by a 405 nm laser and a tunable white-light laser spanning 485–790 nm. Fluorescence emission was detected using two Power HyD detectors for simultaneous multichannel acquisition. Images were acquired using a 20x/0.75 numerical aperture objective and analyzed using Fiji. 20 μL samples were prepared in different ammonium acetate concentrations (100 mM, 25 mM, or 2.5 mM) at pH 8 containing 15 μM tau (+1% Atto655-labelled tau) with and without different RNA variants (supplemented with 2% Cy3-labelled RNA of the same type). Charge matching concentrations were as follows: 0.1 μM for poly-A polydisperse RNA, 0.6 μM for PolyA50, 1.2 μM for PolyA25 and excess negative charge concentrations were 5 μM for all RNA varieties

### Mass photometry

MP experiments were performed using a Refeyn TwoMP mass photometer (Refeyn). To allow RNA size determination, MassGlass UC slides (Refeyn) were treated with poly L-lysine. Therefore, slides were first alternatingly washed with MilliQ water and isopropanol five times (starting and ending with MilliQ washes) before being dried with compressed air. 7 μL of a 0.01% poly L-lysine (Sigma-Aldrich P4707) solution was pipetted onto the middle of a slide, then another washed slide was placed perpendicularly on top, evenly spreading and sandwiching the poly L-lysine between the slides. This assembly was incubated for 30 seconds and a final MilliQ water wash was performed before drying with compressed air.

NT*-GSGAK was buffer-exchanged into 20 mM Tris pH 8.0 using 7K MWCO Zeba™ Spin Desalting Columns (ThermoFisher Scientific). Samples were first pre-diluted into 30 mM ammonium acetate pH 8.0 to concentrations of 20 µM NT*-GSGAK, with and without 50-nucleotide PolyA-RNA (10 µM), and incubated overnight at RT. Immediately prior to measurement, the samples were further diluted in 30 mM ammonium acetate pH 8.0, and final concentrations of 20 nM protein and 10 nM RNA were used on the mass photometry slide. Each measurement was recorded for 3 minutes (9000 frames) to maximise counts with AcquireMP (2025 R1.2) software (Refeyn).

For MP measurements samples containing 15 μM Tau, without and with RNA were prepared in 2.5, 25, and 100 mM ammonium acetate at pH 8 in RNAse free Eppendorf tubes. Samples were further diluted to final tau concentrations of 90 nM in the samples’ corresponding ammonium acetate buffers using RNAse free Eppendorf tubes. Measurements were performed by pipetting a 10-18 μL droplet of the samples’ corresponding ammonium acetate buffer on the glass slide and focusing the laser using the droplet dilution mode in the AcquireMP software (2025 R1.2). Next 2-10 μL (to a total volume of 20 µl) of the prediluted (90 nM Tau) sample was resuspended in the applied droplet and a 1 minute measurement recorded. The ratio of buffer to sample in each recording was adjusted so that the observed total particle counts were between 1000-3000 within a 1 minute recording. The mass photometer was calibrated using the MassFerence™ P1 protein calibrant (Refeyn), with expected mass peaks at 86, 172, 258, and 344 kDa or a RNA ladder with expected mass peaks at 68, 170, 340, and 510 kDa. Mass photometry data were analyzed using DiscoverMP (2025 R1.0) software (Refeyn).

### Native mass spectrometry

For MS experiments NT*-GSGAK was buffer exchanged into 30 mM ammonium acetate pH 8 using 7K MWCO Zeba™ Spin Desalting Columns (ThermoFisher Scientific) following the manufacturer instructions. RNA stocks were diluted as described above in MilliQ water. Final samples contained 15 µM NT*-GSGAK and 7.5 µM 50-nucleotide polyA-RNA and were incubated 72 h at RT. Tau was buffer-exchanged into 25 mM ammonium acetate pH 8 and mixed with RNA to final concentrations of 15 μM Tau and 5 μM 25 nt Poly-A RNA or 50 nt Poly-A RNA. Mass spectra were recorded using a Q Exactive Plus mass spectrometer (ThermoFisher Scientific) modified for high-mass analysis (MS Vision, NL). Samples were loaded into nano-electrospray capillaries (ThermoFisher Scientific, ES380) and analysis was performed using the Xcalibur (Thermo Fisher Scientific) software. The capillary voltage was set to 1.5 kV, the source temperature to 30 °C and the fore vacuum (ion source) was set to 2 mbar. The in-source fragmentation voltage was set to 100 V and the higher-energy collisional dissociation (HCD) voltage was set to 0 V, unless stated otherwise.

### Nuclear magnetic resonance (NMR) spectroscopy

To study the interactions between tau and poly-A polydisperse RNA, 2D NMR^1^H-^15^N-HSQC spectra were recorded. Poly-A polydisperse RNA was titrated upon 500 μL of 128 μM ^15^ N-labelled tau in 20 mM 7.4 pH NaPi buffer with 10% D2O at 298 K, and spectra were measured with a 700 MHz Bruker Avance spectrometer equipped with a cryoprobe. Seven titration steps (0.2, 0.4, 0.7, 1.9, 5, 10, and 42.7 μM) of poly-A polydisperse RNA were used. The spectra from the titration series were compared by determination of the relative intensities (I/I0) based on amplitude intensity, and chemical shift changes (CSC) calculated by Eq.1. Data processing was performed with Bruker TopSpin and analysis in POKY^41^ and RStudio. The cross-peak assignments were based on BMRB ID 50701 and comparison to previous work.^42^

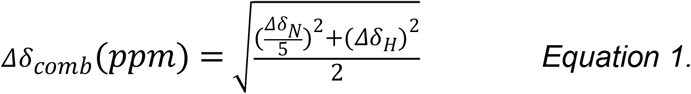

### MD simulations

The initial structure of the 80 residues Tau peptide derivative (residues 281-360) and a single strand 10mer Poly(A) RNA fragment was prepared with the TLEAP module of AMBER 24^43^ using ff19SB^44^ and OL3^45^ force-field parameters for the peptide and RNA respectively. It was placed in the centre of a truncated octahedral box such that the minimum distance between any atom and the box boundaries was at-least 6 Å. The amino- and carboxy-terminus of the Tau peptide was acetylated and amidated respectively. The force-field parameters of ammonium and acetate ions were derived using the GAFF2^46^ force-field and bcc charges through the ANTECHAMBER module of AMBER 24. The net charge of ammonium and acetate was set to “+1e”, and “-1e” respectively. The appropriate number of ammonium and acetate ions equivalent to 25 mM (n=7) salt concentrations was computed based on the volume of the primary simulation box (434115 Å^3^). Packmol^47^ was used to generate the initial distributions of the ammonium and acetate ions within a sphere of radius 20 Å from the centre-of-mass of the Tau peptide+RNA system. Solvation was performed with the OPC^48^ water model and the net charge of the system “-2e” was neutralized by adding two ammonium counterions. MD simulations were performed using the PMEMD module of AMBER 24. The system was initially relaxed through energy minimization with steepest descent and conjugate gradient algorithms. They were then heated to 300 K over 30 ps and equilibrated for 200 ps under NVT and NPT ensembles respectively. Production dynamics was run with NPT conditions for 1 µs. The regulation of the simulation temperature (300 K) and pressure (1 atm), computation of electrostatic interactions, treatment of hydrogen containing bonds and integration of the equation of motion were implemented as described in Lama *et al*.^49^

## Author contributions

AL designed the study with ML. AL, JS, IMP, and HO produced protein. QYN produced RNA and peptides. JS, IMP, QYN, HO, and PM performed and analyzed MP measurements. JS and CM performed and analyzed NMR experiments under supervision of AA. DL performed MD simulations. JS and IMP recorded MS data. MH and AM provided reagents and expertise. AL and ML wrote the paper with input from all authors.

## Acknowledgements

ML is supported by a Karolinska Faculty Career Position, the Swedish Cancer Research Society (Cancerfonden), the Swedish Research Council (VR), the Swedish Society for Medical Research (SSMF) and the Knut och Alice Wallenberg Foundation (KAW). AA is supported by a Consolidator Grant from Karolinska Institutet, the Swedish Research Council, the Swedish Alzheimer Foundation, the Swedish Brain Foundation and CIMED. CM is supported by the Swedish Brain Foundation, Petrus and Augusta Hedlund Foundation, Astrid and David Hageléns Foundation, and the Åhléns Foundation.

