## Supplementary Figures for "Single-molecule mass measurements uncover shifting RNA interactions during condensate phase transitions"

**Table S1.** Sequences and molecular weights of designed proteins

| Name | Sequence | Mass after TEV cleavage (Da) |
|---|---|---|
| NT*-(GSGAE)$_{14}$ | MHTTPWTNPGLAENFMNSFMQGLSSMPG FTASQLDKMSTIAQSMVQSIQSLAAQGRTS PNDLQALNMAFASSMAEIAASEEGGGSLST KTSSIASAMSNAFLQTTGVVNQPFINEITQL VSMFAQAGMNDVSAGSGAEGSGAEGSGA EGSGAEGSGAEGSGAEGSGAEGSGAEGS GAEGSGAEGSGAEGSGAEGSGAEGSGAE ENLYFQSGHHHHHH | 20322.77 |
| NT*-(GSGAP)$_{14}$ | MHTTPWTNPGLAENFMNSFMQGLSSMPG FTASQLDKMSTIAQSMVQSIQSLAAQGRTS PNDLQALNMAFASSMAEIAASEEGGGSLST KTSSIASAMSNAFLQTTGVVNQPFINEITQL VSMFAQAGMNDVSAGSGAPGSGAPGSGA PGSGAPGSGAPGSGAPGSGAPGSGAPGS GAPGSGAPGSGAPGSGAPGSGAPGSGAP ENLYFQSGHHHHHH | 19874.79 |
| NT*-(GSGAK)$_{14}$ | MHTTPWTNPGLAENFMNSFMQGLSSMPG FTASQLDKMSTIAQSMVQSIQSLAAQGRTS PNDLQALNMAFASSMAEIAASEEGGGSLST KTSSIASAMSNAFLQTTGVVNQPFINEITQL VSMFAQAGMNDVSAGSGAKGSGAKGSGA KGSGAKGSGAKGSGAKGSGAKGSGAKGS GAKGSGAKGSGAKGSGAKGSGAKGSGAK ENLYFQSGHHHHHH | 20309.59 |

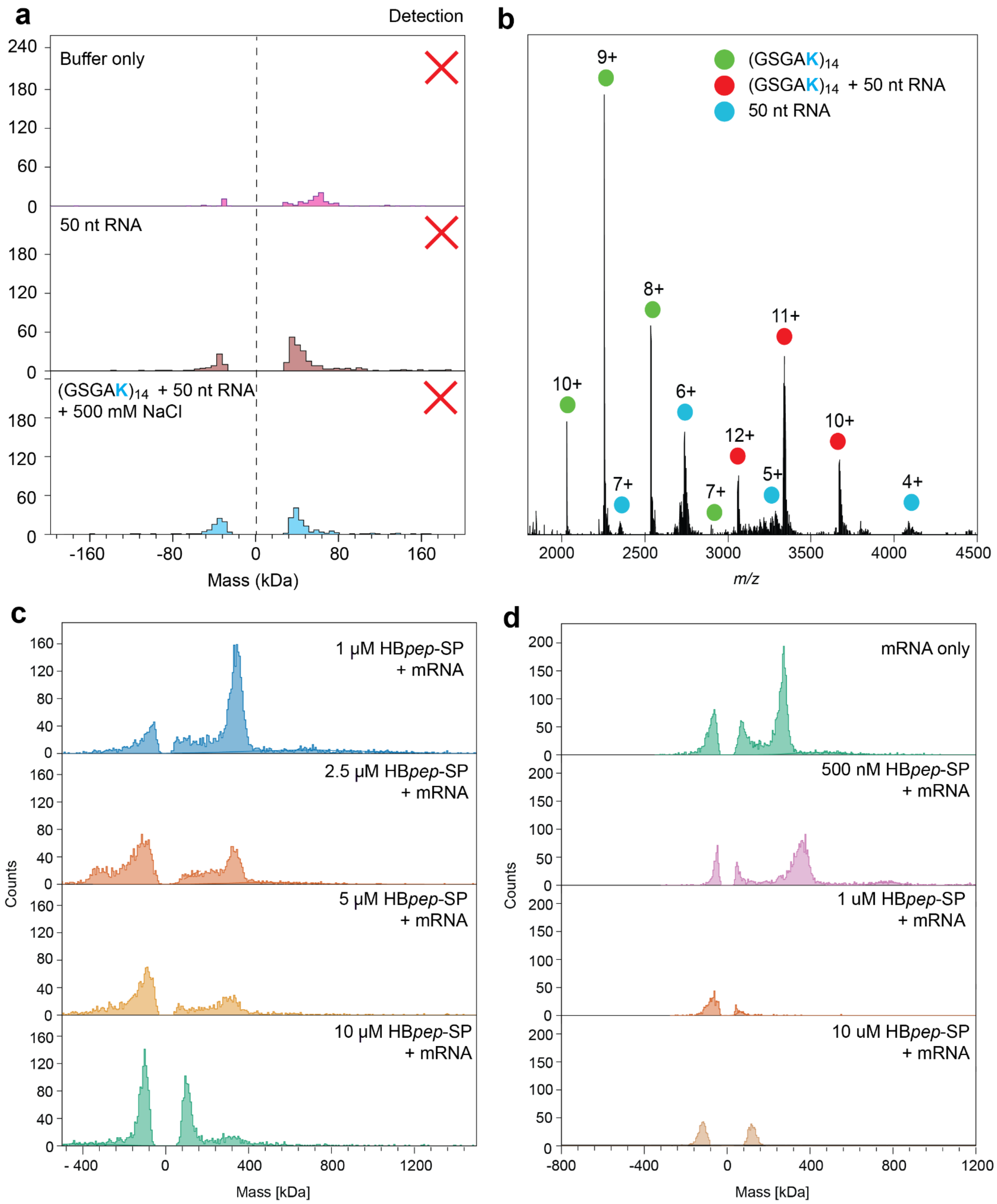


***Figure S1. Buffer-only MP, ratios of unbinding to total signal, and native MS of NT*-GSGAK with 50nt RNA** (a) MP histograms recorded with only buffer, or only 50 nucleotide polyA-RNA show no signal above the anti-binding peak.The signal is suppressed in the presence of 500 mM NaCl. (b) Native mass spectrum of (GSGAK)$_{14}$ incubated with a 3-fold charge excess of 50 nucleotide polyA-RNA shows pronounced 1:1 complexes between protein and RNA. (c) and (d) complete mass histograms for mRNA titrated with HBpep-SP at pH 7.5 (panel c) and pH 5 (panel d).*

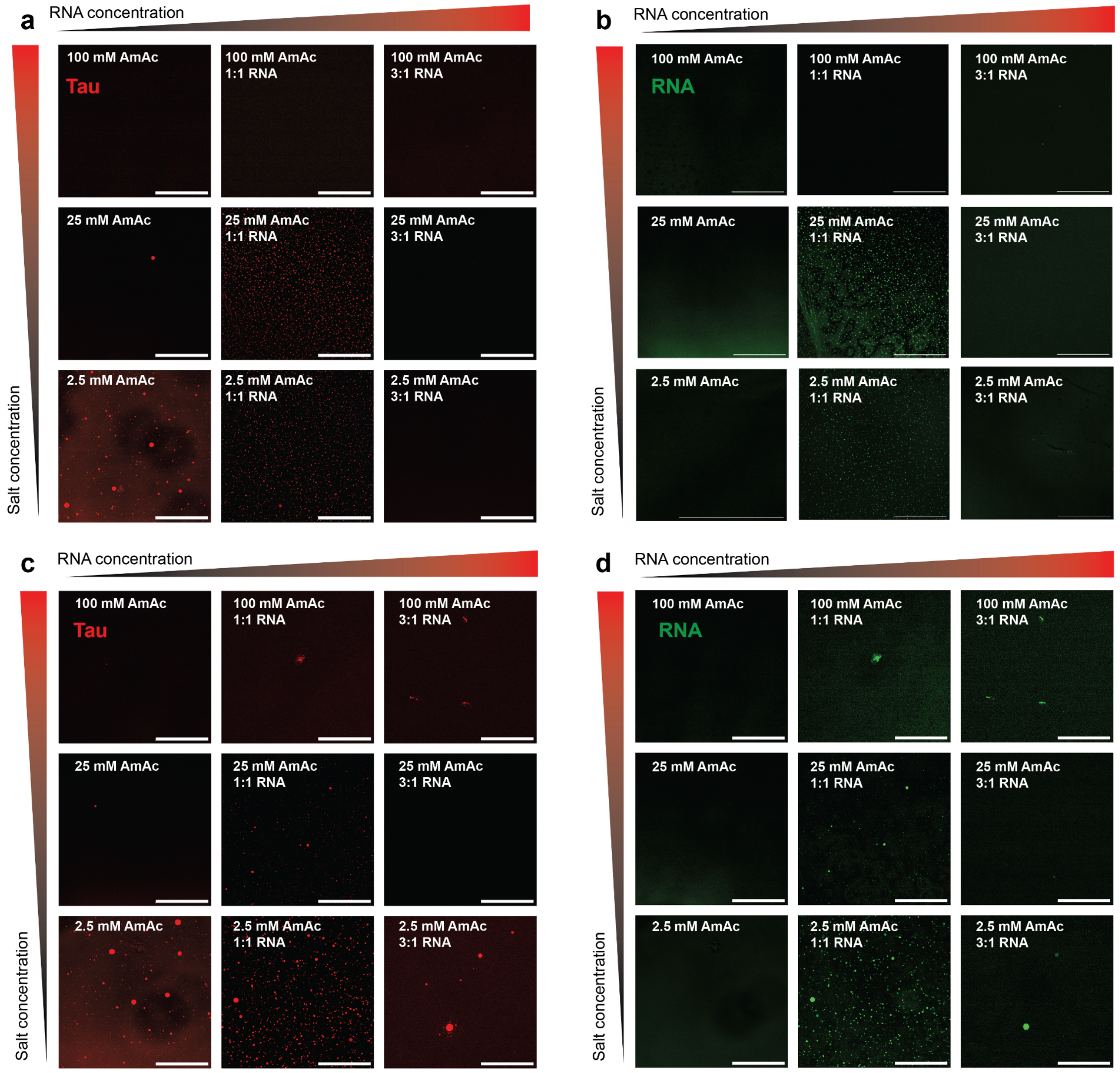


***Figure S2. Microscopy of tau and RNA phase separation at different charge ratios and salt concentrations.*** *(a, b) 15 µM tau without RNA (left), with polydisperse poly-A RNA at a net charge ratio of 1:1 (100 nM RNA with an average length of 300 nucleotides, middle), and at a molar ratio of 1:3 (5 µM RNA, right). AmAc concentrations are 100 mM (top), 25 mM (middle) and 2.5 mM (bottom). Tau is shown in red (a) and RNA in green (b). (c, d) Same as in (a) and (b) with 50-nucleotide poly-A RNA. Scale bars are 200 µm.*

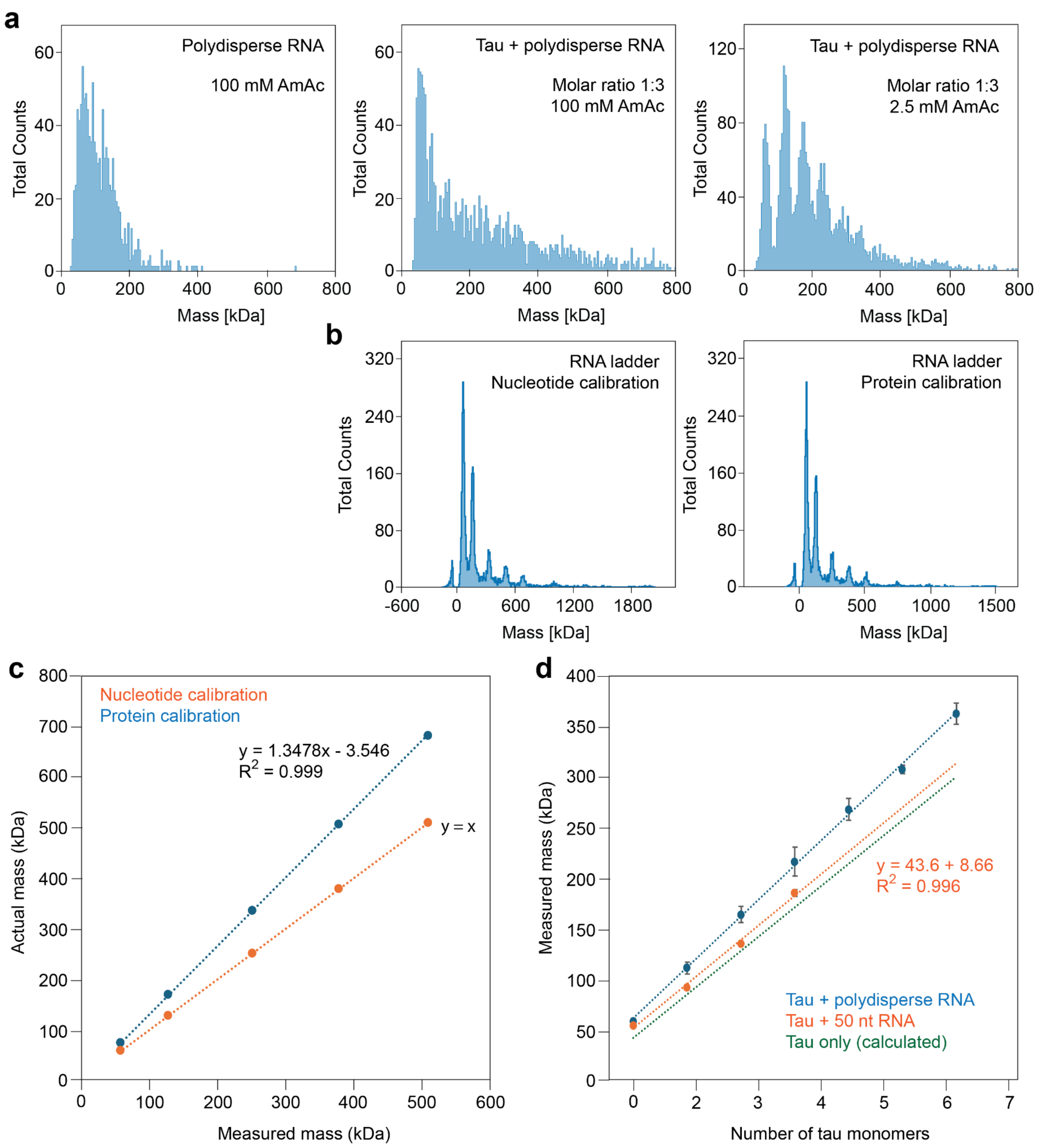


***Figure S3. MP histograms and calibration of tau-RNA complexes.*** *(a) MP histograms of poly-A RNA in 100 mM AmAc (left), tau and polydisperse poly-A RNA at a charge ratio of 3:1 in 100 mM AmAc (middle) and tau and polydisperse poly-A RNA at a molar ratio of 3:1 in 2.5 mM AmAc AmAc (right). No tau complexes are detected in 100 mM AmAc, whereas pronounced complexes can be observed in 2.5 mM AmAc. (b) Histograms of the RNA ladder with nucleotide calibration (left) and protein calibration (right). Note the change in mass axis scale. (c) Plotting the calibrant measured vs. actual masses yields a correction factor of 1.34 when determining RNA masses with a protein calibration. (d) Linear fit of the masses of the tau oligomers formed with 50 nt RNA (orange) show an offset of ca 8.7 kDa (using the protein calibration) and no additional increase in oligomer mass with increasing number of tau monomers.*

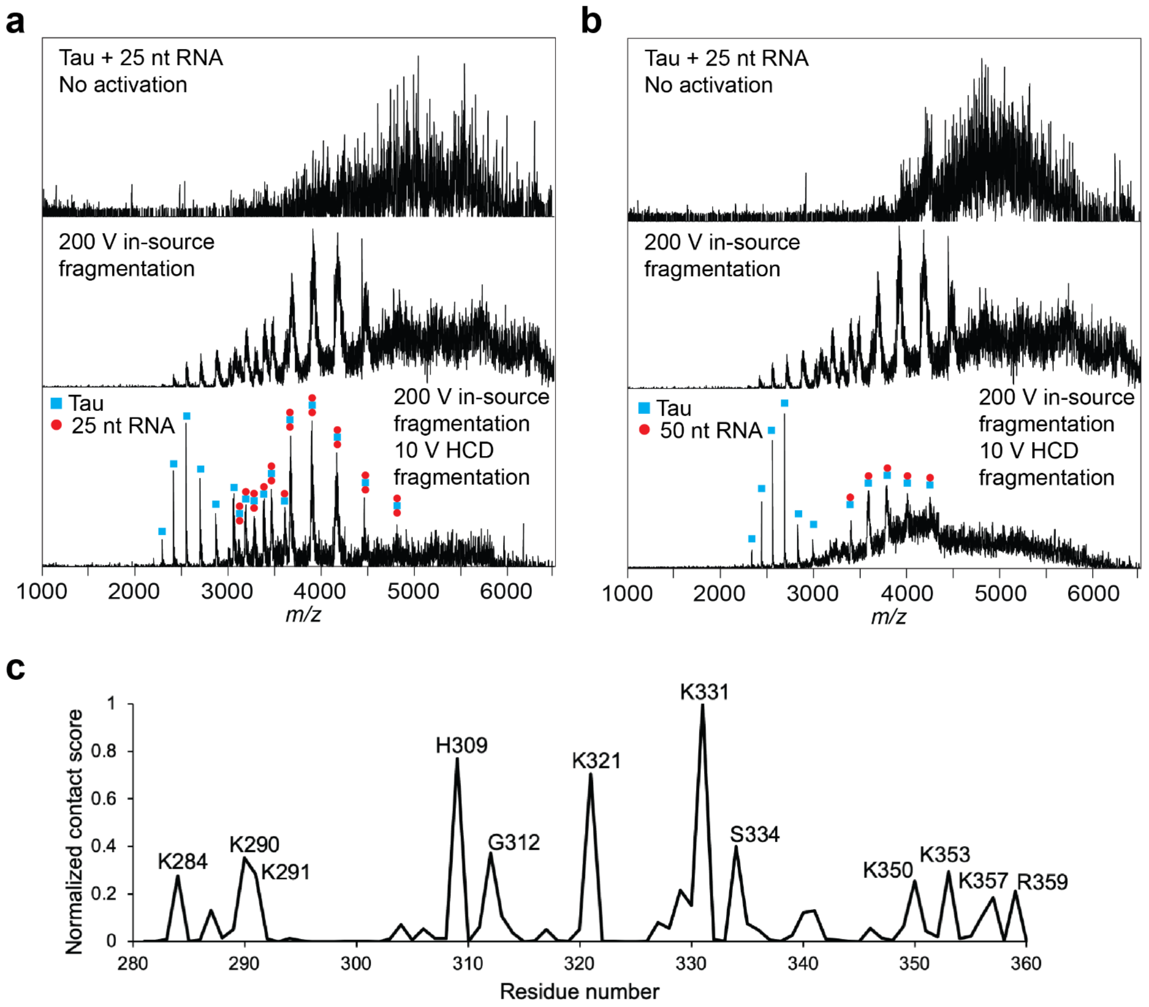


***Figure S4: Native MS and MD of tau-RNA interactions.*** *(a) Native mass spectra of tau in the presence of 25 nt poly-A RNA shows unresolved large assemblies under gentle ionization conditions. Increasing the in-source fragmentation energy releases complexes between one tau and one or two RNA molecules. (b) Native MS of tau with 50 nt poly-A RNA recorded under the same conditions as in (a) yields 1:1 complexes between protein and RNA. (c) Contact scores from all-atom MD simulations of tau repeats 2 and 3 with 30 nt poly-A RNA over the course of a 1 μs simulation shows preferential interactions between RNA and basic residues.*

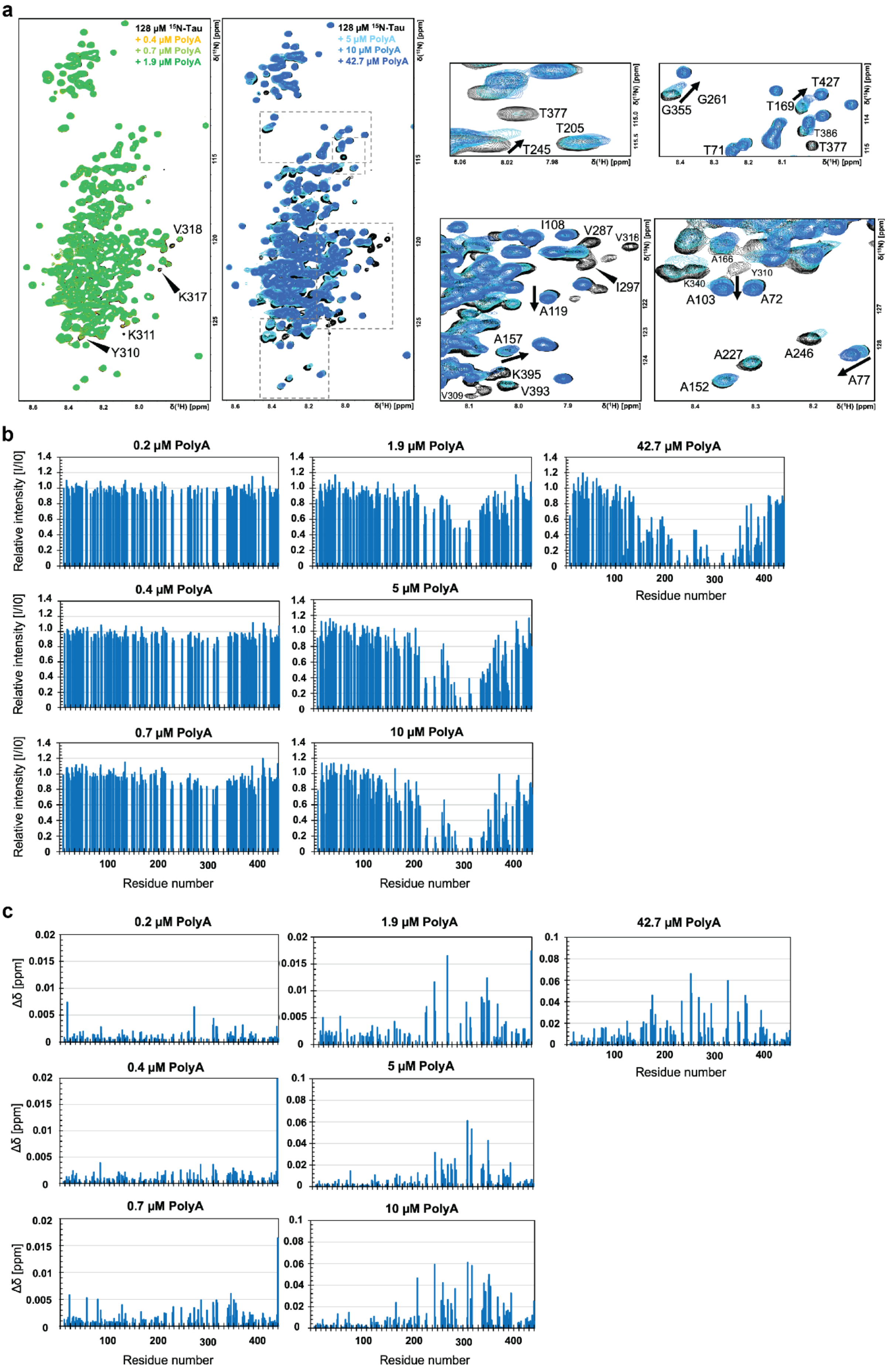
a
128 µM ¹⁵N-Tau
+ 0.4 µM PolyA
+ 0.7 µM PolyA
+ 1.9 µM PolyA
V318
K317
K311
Y310
128 µM ¹⁵N-Tau
+ 5 µM PolyA
+ 10 µM PolyA
+ 42.7 µM PolyA
T377
T205
T245
G355
G261
T427
T169
T386
T377
T71
I108
V287
V318
I297
A119
A157
K395
V393
V309
A166
Y310
K340
A103
A72
A227
A246
A152
A77
δ(¹H) [ppm]
δ(¹⁵N) [ppm]
b
0.2 µM PolyA
1.9 µM PolyA
42.7 µM PolyA
0.4 µM PolyA
5 µM PolyA
0.7 µM PolyA
10 µM PolyA
Relative intensity [I/I0]
Residue number
c
0.2 µM PolyA
1.9 µM PolyA
42.7 µM PolyA
0.4 µM PolyA
5 µM PolyA
0.7 µM PolyA
10 µM PolyA
Δδ [ppm]
Residue number

***Figure S5. NMR spectroscopy data of tau and polydisperse poly-A RNA interactions.*** *(a) 2D NMR $^{1}H$-$^{15}N$-HSQC spectra of 128 mM $^{15}N$-labelled $tau_{441}$ in 20 mM NaP buffer pH 7.4 recorded at 298 K. Titration steps of polydisperse poly-A RNA onto tau are presented as overlayed spectra. (b) The relative intensities for each titration step are determined as $I/I_0$ from the cross-peak amplitude intensities from the spectra in (a). $I_0$ is the cross-peak intensity for tau in the absence of polydisperse poly-A RNA (black spectrum). (c) Changes in cross-peak positions are presented as chemical shift changes for each titration step and calculated using Eq.1.*
